# Social history and exposure to pathogen signals modulate social status effects on gene regulation in rhesus macaques

**DOI:** 10.1101/552356

**Authors:** Joaquin Sanz, Paul L. Maurizio, Noah Snyder-Mackler, Noah D. Simons, Tawni Voyles, Jordan Kohn, Vasiliki Michopoulos, Mark Wilson, Jenny Tung, Luis B. Barreiro

**Author notes:** Department of Psychology, University of Washington, Seattle, WA 98195, USA. ^2^To whom correspondence may be addressed. or. J.T and L.B.B contributed equally to this work.

## Abstract

Social experiences are an important predictor of disease susceptibility and survival in humans and other social mammals. Chronic social stress is thought to generate a pro-inflammatory state characterized by elevated antibacterial defenses and reduced investment in antiviral defense. Here, we manipulated long-term social status in female rhesus macaques to show that social subordination alters the gene expression response to *ex vivo* bacterial and viral challenge. As predicted by current models, bacterial lipopolysaccharide polarizes the immune response such that low status corresponds to higher expression of genes in NF-κB-dependent pro-inflammatory pathways and lower expression of genes involved in the antiviral response and type I interferon (IFN) signaling. Counter to predictions, however, low status drives more exaggerated expression of both NF-κB and IFN-associated genes after cells are exposed to the viral mimic Gardiquimod. Status-driven gene expression patterns are not only linked to social status at the time of sampling, but also to social history (i.e., past social status), especially in unstimulated cells. However, for a subset of genes, we observed interaction effects in which females who fell in rank were more strongly affected by current social status than those who climbed the social hierarchy. Together, our results indicate that the effects of social status on immune cell gene expression depend on pathogen exposure, pathogen type, and social history – in support of social experience-mediated biological embedding in adulthood, even in the conventionally memory-less innate immune system.

## INTRODUCTION

The social environment, both in early life and in adulthood, has a profound and often long-lasting impact on health and mortality in humans and other social mammals (1–5). This relationship is thought to arise in part through changes in gene regulation, which mediate the genomic response to physiological signals of social stress (e.g., glucocorticoids, adrenaline, noradrenaline; (6, 7)). Gene expression signatures of social status and social adversity have now been reported in multiple studies, encompassing clinical and population-based samples in humans, and studies of both experimental and natural populations in other social animals (6, 8–19) (see also (20–25) for evidence in social insects and other social vertebrates). Because this work has concentrated most extensively on peripheral white blood cells, it provides a direct window into how social experiences are reflected in the regulation of the immune system (26–28).

Several broad patterns have emerged from these studies. First, high social adversity, including social isolation, early life insults, and low social status in adulthood, tends to predict higher expression of genes in pro-inflammatory pathways. This observation dovetails with associations between chronic social stress and elevated levels of protein biomarkers of inflammation, including those produced by peripheral blood cells (e.g., interleukin-6; (13, 29, 30)). Second, high social adversity tends to predict lower expression of genes that function in the innate immune defense against virus, especially genes involved in type I interferon (IFN) signaling. In most cases, this pattern has been shown based on data in unstimulated cells (6); however, in bacterial lipopolysaccharide (LPS)-stimulated cells, induction of type I IFN-associated gene expression responses are also attenuated in low status (compared to high status) rhesus macaques (18). Third, these findings are explained in part by socially patterned differences in the use of immune defense-modulating transcription factors. For example, genes that are more highly expressed in low status female rhesus macaques are enriched near accessible binding sites for the transcription factor complex nuclear factor kappa-B (NF-κB), a master regulator of inflammation. In contrast, genes that are more highly expressed in high status females fall near accessible binding sites for interferon regulatory factors, which coordinate type I IFN-mediated responses (18, 31).

These observations have led some authors to propose social environment-mediated trade-offs between antibacterial defense, associated with NF-κB-driven pro-inflammatory signaling, and antiviral defense, associated with the type I IFN response (6, 32). Such a model argues that high social adversity shifts investment in the immune system towards resistance against bacterial pathogens and wound healing (possibly in anticipation of physical insults), at the cost of increased susceptibility to viral pathogens. This in turn accounts for both social gradients in conditions linked to chronic inflammation, such as cardiovascular disease, and social gradients in viral infections. Consistent with this hypothesis, psychosocial stress predicts reactivation rates of latent herpesvirus in mice and humans (33, 34) and rates of respiratory virus infection for experimentally exposed human subjects (35, 36). Similarly, low status cynomolgus macaques males housed in a controlled environment experience both increased susceptibility to experimentally administered adenovirus and elevated rates of coronary artery stenosis (35, 37, 38).

Nevertheless, it remains unclear whether social patterning of viral susceptibility is directly related to social patterning of gene expression in peripheral blood cells. In particular, increased susceptibility to virus has been suggested to occur because the trade-offs induced by social adversity lead to insufficient production of antiviral gene transcripts (32). This logic is largely based on lower expression levels for key antiviral genes (e.g., *MX1, OAS* family genes, interferon regulatory factors) measured at baseline. However, exposure to immune stimulants can radically change the transcriptional landscape of immune cells, and social environmental effects on gene expression have been shown to specifically depend on the cellular environment. For example, in rhesus macaque females, social status has more pronounced effects on gene expression after exposure to LPS, but weaker effects on gene expression after exposure to glucocorticoids (18, 31). Because no study to date has evaluated the effects of social adversity on gene expression after both bacterial and viral challenge, it is therefore unclear whether chronic social stress in fact attenuates the gene regulatory response to virus, consistent with a trade-offs model. In addition, although increased expression of inflammation-related genes and decreased expression of type I IFN-related genes have been associated with social adversity in both adulthood and early life (9, 13, 18, 39) but see (40), we do not yet understand how the timing of social experiences affects the response to either pathogen type.

To address these gaps, we turned to an animal model for social subordination-induced chronic stress: dominance rank in female rhesus macaques. This model takes advantage of the highly hierarchical social structure of female macaques, in which low rank predicts increased harassment and reduced social control, and combines it with the ability to manipulate rank via controlled introduction into newly formed social groups (earlier introduced females are higher ranking) (41, 42). Using this study design, we previously showed that social status effects on blood cell gene expression are pervasive, cell type-specific, and dependent on the cellular environment: most relevant to this study, they can be substantially altered by bacterial stimulation (18). Importantly, we also demonstrated that serial manipulations of dominance rank are possible. Specifically, by rearranging group composition a second time and placing females of previously similar rank into the same social group, the same individuals can be observed when they occupy two distinct positions in the social status hierarchy. Because pre- and post-rearrangement social status are completely uncorrelated, this approach provides an ideal setting to investigate the relative contributions of past versus present social environment to immune gene regulation.

To do so here, we measured genome-wide gene expression levels in 45 adult female rhesus macaques: members of nine, five-member social groups that had undergone two serial rank manipulations spaced ~1 year apart (18, 43, 44). We generated genome-wide expression profiles in blood at baseline, and in blood samples stimulated with bacterial and viral ligands. Finally, we took advantage of historical dominance rank data from the same females to evaluate whether, and for what genes and pathways, immune gene regulation in adulthood is influenced by biological embedding—when social experience leads to systematic, stable biological changes with the potential to influence health (45, 46). Together, our analyses provide new insight into how social subordination-induced chronic stress — both past and present — differentially affects the molecular response to distinct pathogen types.

## RESULTS

### Pathogen exposure- and pathogen type-dependent effects of social status on gene expression

We experimentally manipulated the dominance ranks of 45 adult female rhesus macaques by sequentially introducing them into newly constructed social groups of five females each (**Fig. 1*A***, n=9 social groups, *SI Appendix*, Dataset S1), as described in (18). We maintained these groups for ~1 year (February 2013 - March 2014: Phase I). We then rearranged group composition by performing a second series of sequential introductions, in this case designed so that females from the same or adjacent ranks in Phase I were co-housed in their new groups. We followed the rearranged groups for another 10 months (April 2014 - February 2015: Phase II). As expected from this experimental paradigm (42), earlier introduction predicted higher social status in both phases, as measured by a continuous Elo rating score (Pearson’s r between order of introduction and Elo score: Phase I: −0.57, p = 4.1 × 10^−5^; Phase II: −0.68, p =3.3 × 10^−7^). Importantly, once dominance ranks were established, they remained highly stable throughout each study phase, and individual Elo scores were completely uncorrelated between phases (r=0.06, p=0.68, *SI Appendix*, Fig. S1).

**Fig. 1.**
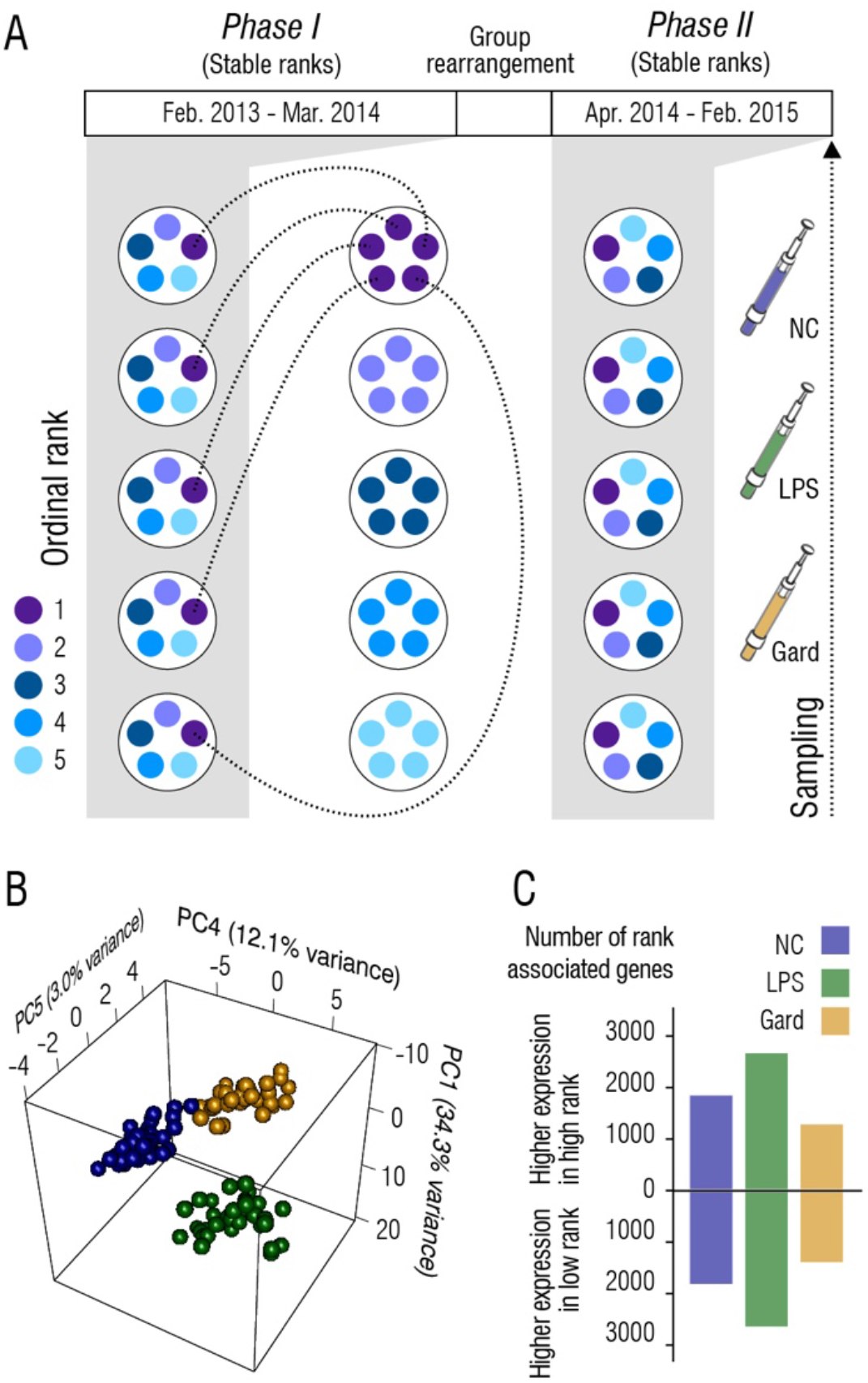
Social status effects on gene expression within and across conditions. *(A)* Schematic representation of group formation in Phase I, mid-study rearrangement, and reorganization into new hierarchies in Phase II. *(B)* Principal components analysis of gene expression data across all three conditions. The combination of PC1, PC4 and PC5 separates unstimulated negative controls (NC, blue) from samples stimulated with either LPS (green) or Gard (yellow). *(C)* Number of rank-associated genes (FDR<5%) that are more highly expressed in high ranking females (top bars) or low ranking females (bottom bars), within each condition.

To characterize the effects of social status on the immune response to bacterial versus viral challenge, including the contribution of social history, we obtained blood samples from each study subject in Phase II of the study. We generated three gene expression profiles per animal, from: (i) a control (untreated) sample, with blood cultured in cell culture media only (negative control, NC); (ii) a sample cultured in cell culture media spiked with LPS, a Toll-like receptor 4 (TLR4) agonist that mimics infection by Gram-negative bacteria; and (iii) a sample cultured in cell culture media spiked with Gardiquimod (Gard), a Toll-like receptor 7 (TLR7) agonist that mimics infection by a single-stranded RNA virus. We incubated samples from each individual for 4 hours in parallel, lysed the white blood cell fraction, and generated RNA-sequencing data from the matched non-stimulated and stimulated conditions. To confirm successful immune stimulation, we performed a principal component analysis (PCA) on the correlation matrix of normalized gene expression levels for all conditions, after controlling for relatedness, age, and the potential confounding effects of batch and tissue composition (**Fig. 1*B***; *SI Appendix*, Material and Methods). We observed distinct clusters corresponding to the control, LPS-stimulated, and Gard-stimulated samples, with treatment effects most clearly reflected along PC1 (r=0.61, p=6.5 × 10^−10^ for the separation between control versus LPS samples) and PC4 (r=0.76, p=1.7 × 10^−17^ for the separation between control versus Gard samples). As expected, genes up-regulated after stimulation were significantly enriched for immunological and inflammatory processes canonically associated with bacterial and viral defense (see *SI Appendix*, Dataset S2).

Consistent with previous findings (18, 19, 34), we also observed a strong signature of dominance rank. Specifically, dominance rank (Elo score) was significantly correlated with PC3 of the full gene expression matrix within all three conditions (Pearson’s r = 0.75 (NC); r = 0.75 (LPS), r = 0.69 (Gard), p<6.0 × 10^−7^ for all conditions; *SI Appendix*, Fig. S2). This observation translated to gene-level analyses, where dominance rank drove the expression of 3,675, 5,322 and 2,694 genes (FDR<5%) in the control, LPS, and Gard conditions, respectively (**Fig. 1*C***, *SI Appendix*, Dataset S3). Strikingly, the number of rank-associated genes in the LPS condition was 1.5 – 2x higher than in the Gard or control conditions. Thus, although rank effects are amplified after activation of the bacterial-sensing TLR4 pathway, this pattern does not appear to be a universal feature of immune activation. Indeed, the number of genes for which the intensity of the response (i.e., gene expression in LPS/Gard conditions, relative to paired control samples) depended on dominance rank was almost fivefold lower in the viral Gard condition than for the bacterial LPS-stimulated samples (851 vs 4,111; FDR < 0.05; **Fig. 2*A***). Further, while 73% of rank effects on the LPS response were directionally biased towards larger responses in low status females, neither low nor high status systematically predicted stronger responses to Gard. Instead, rank effects on the response to Gard were almost perfectly balanced: 49.5% of rank-dependent genes exhibited a stronger response in low status animals, while 51.5% exhibited a stronger response in high status animals (**Fig. 2*B***).

**Fig. 2.**
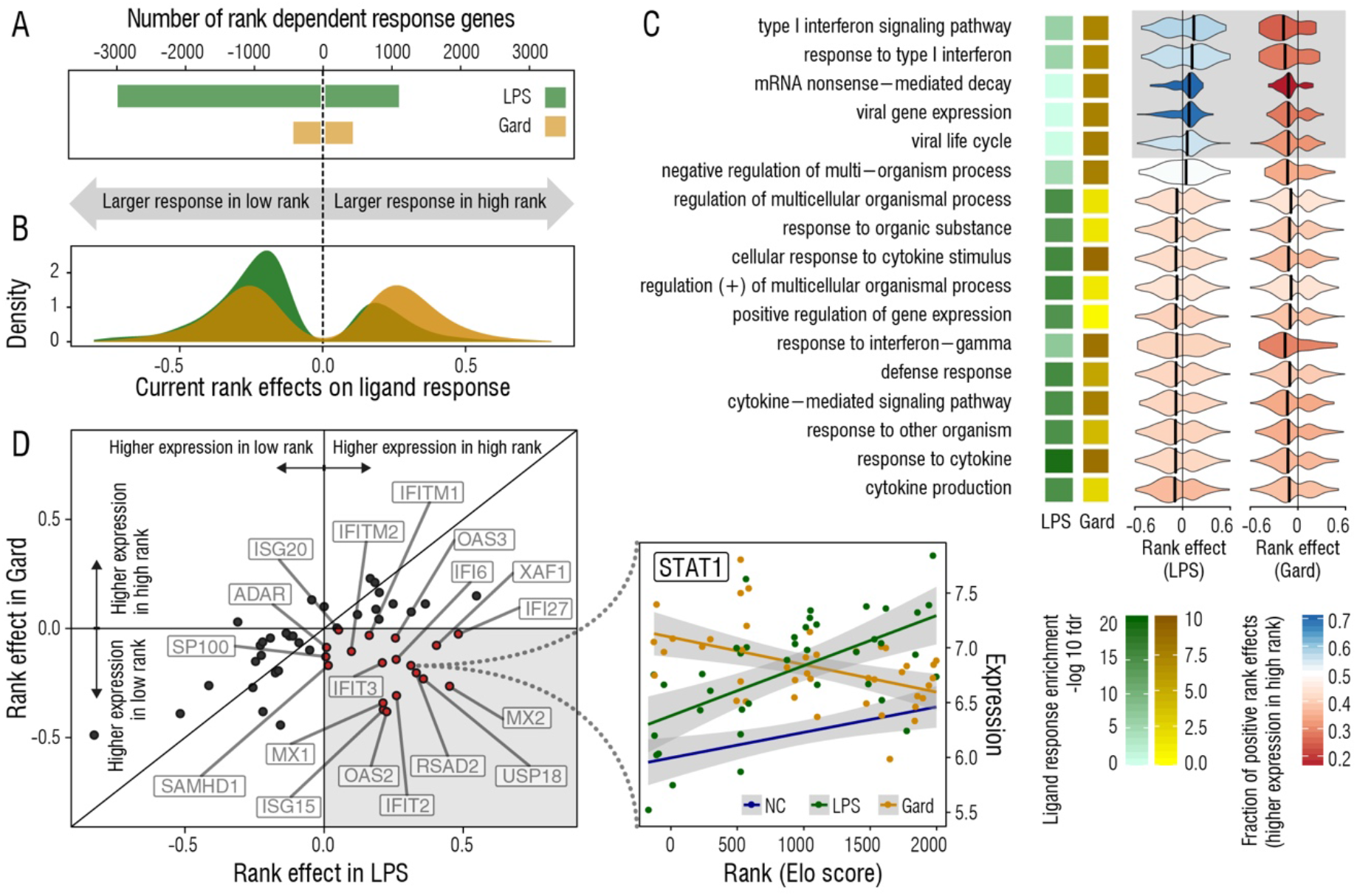
Contrasting effects of dominance rank in cells challenged with a bacterial versus viral mimic. *(A)* More genes show a rank-dependent response (stimulated condition compared to paired control sample) after challenge with LPS than with Gard. LPS challenge also leads to asymmetric responses, in which the response is more commonly stronger in low-ranking females (left bar) than in high-ranking females (right bar); no such asymmetry is observed after Gard challenge. *(B)* Distribution of effect sizes among genes for which the absolute magnitude of the response to LPS (green) and Gard (yellow) depends on dominance rank. After LPS stimulation, most genes respond more strongly in low status females, but rank effects on the response to Gard are more balanced. *(C)* Polarization of rank effects in the union of the top ten Gene Ontology categories, per condition, that were most significantly enriched among genes up-regulated by LPS/Gard stimulation. For each category, darker squares correspond to stronger statistical support for enrichment in the LPS response (green) and Gard response (yellow), respectively, for genes upregulated relative to controls. Horizontal violin plots are colored based on the proportion of rank-associated genes in each category that are more highly expressed in high-ranking individuals, separately for the LPS and Gard conditions. Positive x-axis values: genes that are more highly expressed in high-ranking females; negative x-axis values: genes that are more highly expressed in low-ranking females. The gray shaded box highlights gene categories for which the distributions of rank effects significantly differ between LPS and Gard conditions (Wilcoxon test: Benjamini-Hochberg FDR-corrected p<9×10^−3^). *(D)* Rank effects on gene expression in the LPS (x-axis) versus Gard (y-axis) conditions, for genes in the Gene Ontology category “type I interferon signaling pathway” that were rank-associated in the LPS condition, Gard condition, or both. Labeled genes in the lower right quadrant are cases in which rank effects are directionally reversed in LPS versus Gard-stimulated samples. They include key master regulators of the response to virus such as *STAT1*, which is more highly expressed in high status females in control and LPS conditions, but more highly expressed in low status females in the Gard condition (*STAT1* plot y-axis: normalized, log-transformed gene expression). In *(C)* and *(D)*, all genes affected by rank at an FDR of 20% or less are plotted (the overall pattern is qualitatively unchanged at a more stringent FDR threshold).

To investigate the hypothesis that social status-induced stress mediates trade-offs between antibacterial and antiviral responses, we restricted our analysis to genes belonging to the Gene Ontology categories that were most enriched among genes up-regulated by LPS, Gard, or both. As we showed in previous analyses (18), genes involved in immune defense, inflammation and cytokine signaling are biased towards higher expression in low-ranking females, in LPS condition samples (**Fig. 2*C***). In contrast, genes involved in viral defense-associated type I IFN signaling are biased towards higher expression in high-ranking females, in the same condition (**Fig. 2*C***). A trade-offs model would predict a similar dichotomy after challenge with a viral mimic. Surprisingly, this is not what we observed: in the Gard condition, low status predicted increased expression of both pro-inflammatory/cytokine signaling-associated and type I IFN-associated genes. For example, for rank-associated genes in the Gene Ontology category “type I interferon signaling” 57% of genes measured in the LPS condition were more highly expressed in high status females. However, for rank-associated genes in the same category, measured in the Gard condition, only 25% were more highly expressed in high status females. Consequently, status-related effects for type I IFN genes differ significantly between LPS and Gard conditions (Wilcoxon test, p=6.4 × 10^−4^). This pattern holds for key viral defense genes such as *OAS2* and *OAS3*, which are involved in inhibition of viral replication; the IFN-inducible genes *IFIT2, IFIT3, MX1* and *MX2;* and *STAT1*, a master regulator of interferon-mediated defense (**Fig. 2*D***). It also extends to measures of the response to immune challenge (i.e., the change in gene expression levels between control and challenged cells): genes involved in type I IFN signaling tend to be more strongly up-regulated in high status females after LPS challenge, but more strongly up-regulated in low status females after Gard challenge (*SI Appendix*, Fig. S3)

Previous studies suggest that social environmental effects on gene expression arise through socially structured differences in immune defense-associated transcription factor (TF) binding (e.g., (10, 13, 18)). We therefore investigated the TFs that might account for rank effects on gene expression during the immune response to bacteria (LPS condition) or virus (Gard condition). To do so, we identified predicted TF binding sites in gene promoter regions that are also accessible to TF binding (i.e., in open chromatin regions identified using previously collected ATAC-seq data from rhesus macaque peripheral blood mononuclear cells (18)). As previously reported (18), genes that were more highly expressed in low status animals were enriched for NF-κB binding sites in the LPS condition. In contrast, genes that were more highly expressed in high status animals were enriched for predicted TF binding sites for interferon regulatory factors (IRF1, IRF2, and IRF7) and STAT1— the master-regulators of anti-viral responses (**Fig. 3**). Strikingly, that dichotomy disappeared when samples were stimulated with a viral mimic. In the Gard condition, the promoter regions of genes that were more highly expressed in low status animals were enriched for TF binding sites for virtually all immune-associated TFs, including NF-κB, several IRFs, and STAT1 (**Fig. 3**). These results corroborate our enrichment analyses for the gene expression data alone (**Fig. 2C**), and suggest that differences in TF activity account for the distinct patterns of social status-associated gene expression after LPS versus Gard stimulation.

**Fig. 3.**
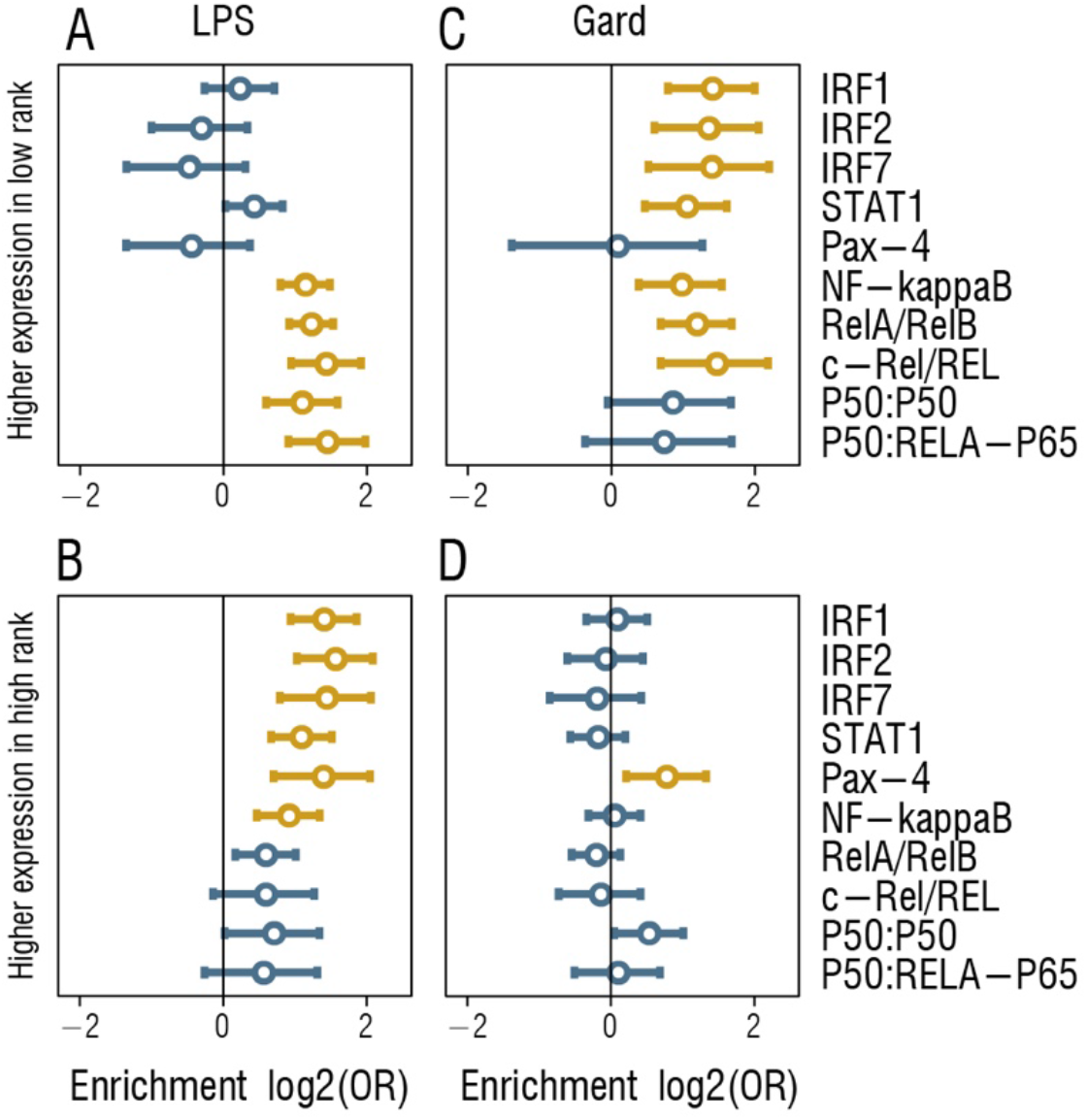
Transcription factor binding sites enriched near rank-associated genes differ between LPS and Gard-challenged cells. In the LPS condition, *(A)* Predicted NF-κB and NF-κB subunit (RelA/RelB/p50/p65) binding sites are enriched in the promoter regions of genes that are upregulated by LPS and more highly expressed in low-ranking females, while *(B)* predicted interferon regulatory factor (IRF) and STAT1 binding sites dominate the enriched categories for genes that are upregulated by LPS but more highly expressed in high-ranking females. This polarization disappears in the Gard condition, where *(C)* Predicted NF-κB, NF-κB subunit, IRF, and STAT1 binding sites are all enriched in the promoter regions of genes that are upregulated by Gard and more highly expressed in low-ranking individuals, with *(D)* no clear signature of immune-related TF binding site enrichment among genes that are upregulated by Gard but more highly expressed in high-ranking individuals. Promoter regions are defined as the 5 kb upstream of each gene transcription start site; in each category, enrichment analyses included all rank-associated genes that passed an FDR cutoff of 20%. Error bars show the 95% confidence interval for the log_2_(odds ratio), and those TFs shown in yellow indicate significant enrichment at an FDR threshold of 10% (p<1.3 × 10^−3^).

### Social history effects on immune gene regulation

Although the gene expression data were generated in Phase II, we also collected behavioral data that allowed us to quantify rank in Phase I, prior to the mid-study rank rearrangement (samples were collected 9.01±0.60 s.d. months after Phase I groups were dissolved and 7.65±0.50 s.d. months after each animal was introduced into her Phase II group). We took advantage of this study design to investigate whether past social status affected immune cell gene expression independently from social status at the time of sampling (“current rank”), in support of biological embedding. Strikingly, we identified a strong global signature of past social status on gene expression levels as well: past Elo score was correlated with PC2 within each condition (control: Pearson’s r = −0.76, p=2.4 × 10^−9^; LPS: r = −0.58, p=8.0 × 10^−5^; Gard: r = −0.44, p=3.8 × 10^−3^; *SI Appendix*, Fig. S2).

On the level of individual genes, social history effects were detectable in all three conditions, but were by far more apparent in the unstimulated, control condition than in the LPS or Gard conditions (**Fig. 4*A***). For example, at an FDR of 10%, we identified 3,735 past rank-associated genes in the control condition, compared to 1,712 and 141 in the LPS and Gard conditions, respectively (**Fig. 4*A***). This pattern is highly robust across statistical thresholds (*SI Appendix*, Fig. S4). Genes associated with past rank were also enriched for distinct biological functions (*SI Appendix*, Dataset S2). For example, in the unstimulated controls, genes that were more highly expressed in high status animals at the time of sampling were strongly enriched for viral transcription (FDR-corrected p=2.8 × 10^−11^) and viral gene expression (FDR-corrected p=5.5 × 10^−11^). However, genes associated with past rank were not overrepresented in either category. Similarly, genes that were more highly expressed in females with a past history of high status were uniquely enriched in categories involved in the epigenetic regulation of gene expression, including chromatin organization (FDR-corrected p=4.8 × 10^−11^), and histone modification (FDR-corrected p=3.6 × 10^−7^) (*SI Appendix*, Fig. S5, Dataset S2).

**Fig. 4.**
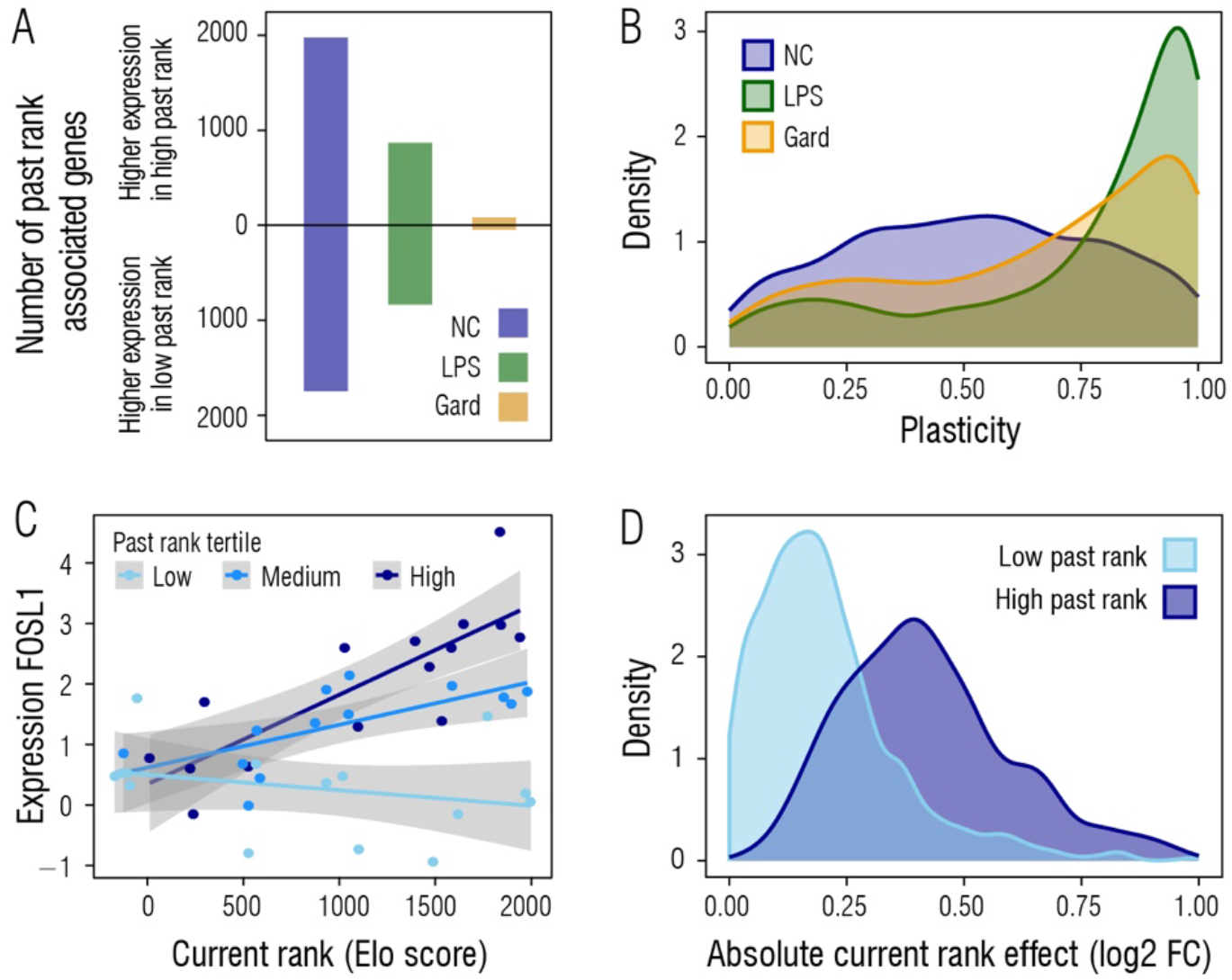
Social history effects on immune gene regulation. *(A)* Number of past rank-associated genes (FDR<10%) that are more highly expressed in high past-ranking females (top bars) or low past-ranking females (bottom bars), within each condition. Past rank effects are more common in the unstimulated control samples than in LPS- or Gard-stimulated samples. *(B)* Distribution of plasticity scores (Θ) for each condition, for the top 1000 rank-associated genes (based on total variance in gene expression explained by the combination of past and current rank: see also *SI Appendix*, Materials and Methods, Fig. S6). Plasticity scores are systematically lower in the unstimulated control samples (NC) than in the LPS or Gard conditions, indicating that immune stimulation proportionally weakens social history effects, relative to current rank effects. *(D)* In an example of a current rank x past rank interaction, current rank effects (x-axis) on expression of the transcription factor *FOSL1*, which is implicated in regulation of the type I IFN response (47), are strongest in females who were previously high rank and weakest in females who were previously low rank (_interaction_=0.54, FDR-corrected p=0.01). *(E)* Predicted current rank effects, based on model fits for each gene on the full data set (see *SI Appendix*, Materials and Methods), for low past rank and high past rank females (based on the mean Elo scores for the lowest ranking and highest ranking females in Phase I groups, respectively). Distributions show estimated effect sizes across 1,079 genes for which we identified a significant current rank-past rank interaction effect in the control condition. Current rank effects were systematically larger in females who were previously high-ranking than in females who were previously low-ranking (Wilcoxon test p<2.2×10^−16^).

To investigate the relative contribution of social history versus current rank at a more granular scale, we developed a gene-specific measure of plasticity (Θ) in each condition. We defined Θ as the square root of the ratio between the variance in gene expression explained by current rank, and the total variance explained by both past and current rank (*SI Appendix*, Materials and Methods). Θ values range from 0 to 1, where values close to 1 imply a high degree of plasticity and little evidence of memory, and values close to 0 imply a high degree of memory and little plasticity in response to changes in social status. Overall, we identified a much greater contribution of social history to gene expression levels in the unstimulated control samples than in either of the stimulated conditions (control: median Θ for the top 1000 rank-associated genes in NC = 0.52; LPS = 0.87; Gard = 0.74; Wilcoxon test p<2.2 × 10^−16^ for all pairwise comparisons; **Fig 4*B***; see also *SI Appendix*, Fig. S6 for comparisons using different numbers of rank-associated genes). In agreement with this observation, social status effects on the response to LPS and Gard (i.e., the interaction between Elo score and control versus stimulated treatment) were dominated by the effects of current rank. We identified 4,111 and 851 genes for which the response to LPS and Gard, respectively, depended on dominance rank at the time of sampling (FDR<5%), but no cases in which these responses depended on past rank. Thus, although there is a strong signature of social history on gene expression levels in an unperturbed state, this signature gives way to the effects of current social conditions when cells enter an immune-challenged state.

Finally, females with the same social status in Phase II could come from different social histories: some climbed the social hierarchy, some fell, and others maintained a consistent position. We therefore tested if past and current rank combined non-additively to influence gene expression. In the control condition, where the effects of social history are most pronounced, we identified 1,079 genes in which past and current rank interacted to influence gene expression (FDR<5%). Interaction effects were strongly directionally biased: specifically, for 88% of these genes, a history of past low status predicted reduced sensitivity to current dominance rank (**Fig. 4*C* and 4*D***). Thus, females who fell in rank were strongly affected by their current, lower rank in Phase II, whereas females who achieved higher status in Phase II were proportionally more affected by their lower status in Phase I.

## DISCUSSION

Convergent evidence from humans, wild animal populations, and experimental animal models indicates that social interactions are reflected in the regulation and activity of the immune system (6, 7, 11, 26–28, 48). Our findings join those of others to suggest that social adversity is particularly relevant to the inflammatory response, one of the first lines of defense in the innate immune system (49). Because biomarkers of inflammation in turn predict disease and mortality outcomes (50), these findings suggest that social regulation of immune gene expression may partly mediate social gradients in health. However, contrary to current predictions based on data collected in unstimulated conditions, low status females did not mount attenuated gene regulatory responses to viral challenge (6, 32). Instead, they show a stronger up-regulation of genes involved in the regulation of type I IFN when comparing Gard to control samples. Further, for these genes, the status-dependent polarization of gene expression patterns observed following exposure to a bacteria-associated challenge disappears following exposure to a virus-associated challenge.

These results suggest that baseline patterns of immune gene expression provide limited insight into the gene regulatory response to actual immune stimuli. Thus, while the gene expression signature of social stress may be somewhat conserved across different types of social adversity, and potentially across species, it does not appear to be highly conserved across pathogen environments. Our findings are consistent with previous reports that social status interacts with LPS and glucocorticoid exposure (18, 31), and that genetic effects on gene expression can also be altered by local cellular conditions (i.e., gene-environment interactions, e.g., (51–54)). However, while prior work has primarily shown environment or genotype-dependent differences in the presence or magnitude of effects, here we observed a more striking pattern: directional shifts in the effects of social status, specifically for genes involved in the antiviral response. This reversal of effects suggests that social status-sensitive regulatory elements involved in the response to LPS are, at least to some degree, distinct from social status-sensitive regulatory elements involved in the response to Gard — a hypothesis that requires further experimental validation.

Our findings also highlight a novel way in which social environmental effects on gene regulation depend on context. Specifically, we identified thousands of genes for which gene expression levels in peripheral blood cells depend not only on current social status, but also on the effects of past social status. Social history effects were themselves dependent on other factors, however, with much more extensive effects in unstimulated cells than immune-challenged cells, and in females with a history of low status than for females with a history of high status. These observations are consistent with several possibilities. First, a history of high social status (i.e., low social adversity) could confer increased plasticity in response to environmental change (e.g., subsequent losses of status). Second, historical exposure to social subordination-induced stress could blunt responses to future high-quality environments. Third, social history effects may dissipate over time, but at a faster pace for formerly high status females. Given that the effects of current rank dominate after immune stimulation, external environmental challenges might hasten this process. Longitudinal, repeated measures of gene expression will be needed to differentiate between these possibilities.

Regardless of the explanation, our observations clearly highlight the importance of social history to immune gene expression patterns measured many months later. They therefore extend the concept of biological embedding to adulthood, in keeping with the observation that the molecular mechanisms thought to mediate the embedding process early in life (e.g., DNA methylation, histone marks) remain environmentally sensitive across the life course. The mechanism of memory in this case may differ from cases of biological embedding that persist over years or decades. For example, memory T cells, which can represent up to 40% of total circulating T cells in adulthood (55), can survive in the body for years, raising the possibility that slow turnover of some PBMC subsets could be responsible for the social history effects we observed. However, recent evidence suggests that even short-lived cells in the innate immune system—which canonically has been thought to lack memory—can be epigenetically altered by environmental experience (56).

Indeed, there is very strong evidence that both early life social adversity and social adversity in adulthood shape health and fitness outcomes (2, 57, 58). If temporally separate exposures independently tap into the same gene regulatory mechanisms, then the genome can be viewed as a mosaic, in which some loci retain more of a long-term signature of social history than others. This perspective is consistent with studies of organism-level traits. For example, in nonhuman primate models of social stress, behavioral signatures of status are highly plastic with changes in dominance rank, whereas the development of coronary artery plaques are explained by the combination of present and past social status (59, 60). A key contribution of genomic approaches rests in their ability to compare levels of plasticity across thousands of traits simultaneously. Such studies promise to reveal which pathways are more important for encoding memory versus sensing the current environment—an important step towards understanding the molecular mechanisms that affect resilience and recovery.

Finally, our analyses suggest that there is no simple mapping between social environmental effects on immune cell gene expression and their effects on disease and mortality risk. Notably, although the type I IFN response is often discussed in opposition to pro-inflammatory, NF-κB-mediated antibacterial responses, type I IFN signaling itself can also be pro-inflammatory (61). High reactivity to both viral and bacterial ligands may therefore represent two distinct sources of elevated inflammation in low status individuals. Alternatively, because Gardiquimod only models TLR7 stimulation by single-stranded RNA viruses, it remains possible that other types of viral challenges produce responses more consistent with social status-mediated trade-offs. For instance, while many of the viruses that have been linked to social gradients are single-stranded RNA viruses (e.g., rhinovirus, respiratory syncytial virus, coronavirus), others are double-stranded DNA viruses (e.g., Epstein-Barr virus, cytomegalovirus, and other herpesviruses) that activate distinct immune pathways at the cell surface, in endosomes, and in the cytoplasm (e.g., TLR2, TLR9, and AIM2-dependent signaling, respectively) (62).

## MATERIAL AND METHODS

### Study Population

The primary set of study subjects were 45 adult female rhesus macaques (*Macaca mulatta*) housed in demographically uniform social groups of five each (n = 9 social groups) at the Yerkes National Primate Research Center (YNPRC) Field Station. These animals were part of a larger, multi-year study investigating social status effects on behavior, physiology, and gene regulation in immune cells (18, 43, 44, 60). All study subjects were initially drawn from the YNPRC breeding colony, assigned to five-member social groups in January – June 2013, and behaviorally monitored to assess social status in February 2013 - March 2014 (Phase I). Group membership was rearranged in March – June 2014 (start of Phase II) by co-housing females of the same or adjacent ordinal ranks in Phase I in the same Phase II groups. This approach allowed us to observe all possible between-phase changes in ordinal dominance rank and produced completely uncorrelated individual ranks between phases (r=0.06, p=0.68), thus enabling us to separate the effects of rank at sampling from rank social history.

In both phases, social groups were formed by sequentially introducing females into indoor-outdoor run housing (25 m by 25 m for each area) over the course of 2 - 15 weeks. Behavioral data on affiliative and agonistic interactions were collected in both phases using focal sampling (345 total hours of observation, 223.5 hours in Phase I and 121.5 hours in Phase II) (63). Based on these data, dominance rank values were assigned using Elo ratings, a continuous measure of rank in which higher scores correspond to higher status and ratings are updated following any interaction in which a winner or loser can be scored (64, 65). All blood samples for gene expression analysis were collected in Phase II, 7.65±0.50 s.d. months after females were first introduced into their Phase II groups. These rank values were highly stable, with an Elo stability index from 0.995 - 1.00 (n=9 groups), where 1 corresponds to a hierarchy in which higher-ranking females always win competitive encounters with lower-ranking females, with no rank reversals (66).

Information on group membership, age, and dominance rank for all study subjects is provided in *SI Appendix, Dataset S1*.

### Blood Sample Collection and *In Vitro* Challenges

We set out to test how dominance rank affects the immune response to bacterial versus viral stimulation and to investigate social history effects at both baseline and in response to pathogen stimulation. To do so, we drew 1 mL of whole blood from each female into each of three TruCulture blood collection tubes (Myriad RBM) containing either: *(i)* cell culture media only (negative control: “NC”); *(ii)* cell culture media plus 1 μg/mL of *E. coli*-derived lipopolysaccharide (0111:B4 strain: “LPS” condition); or *(iii)* cell culture media plus 1 μg/ml of the TLR7 agonist Gardiquimod (“Gard” condition), a synthetic ligand that mimics infection with a single-stranded RNA virus. Samples were incubated in parallel for 4 hours at 37°C. We then separated the serum and cellular fractions, lysed and discarded the red cells from the cell pellet with red blood cell lysis buffer (RBC lysis solution, 5 Prime Inc.), and lysed the remaining white blood cell fraction in Qiazol for storage at −80°C. We extracted total RNA from each sample using the Qiagen miRNAEASY kit. Previous analyses of the NC and LPS samples are reported in (18). Gard samples were also collected at the same time, but gene expression data for these samples were not generated until a later date (see below for discussion of possible associated batch effects). Importantly, raw data for all three conditions were re-processed and re-normalized in parallel for this analysis.

To control for cellular compositional effects, we also drew an additional blood sample in the same draw to measure the proportional representation of ten white blood cell subsets: classical monocytes (CD14^+^/CD16^−^), CD14^+^ intermediate monocytes (CD14^+^/CD16^+^), CD14^−^ non-classical monocytes (CD14^−^/CD16^+^), helper T cells (CD3^+^/CD4^+^), cytotoxic T cells (CD3^+^/CD8^+^), double positive T cells (CD3^+^/CD4^+^/CD8^+^), CD8^−^ B cells (CD3^−^/CD20^+^/CD8^−^), CD8^+^ B cells (CD3^−^/CD20^+^/CD8^+^), natural killer T lymphocytes (CD3^+^/CD16^+^), and natural killer cells (CD3^−^/CD16^+^). The antibody-fluorophor combinations used for flow cytometry are provided in (18), and estimated cell type proportions for each sample are provided in *SI Appendix, Dataset S1*.

### RNA-Seq library preparation, low-level data processing, and batch effect correction

Each RNA-sequencing library was prepared from 200 ng of total RNA using the NEBNext Poly(A) mRNA Magnetic Isolation Module and the NEBNext Ultra RNA Library Prep Kit (New England Biolabs), following the manufacturer’s instructions and selecting for ~350 bp size fragments. Libraries were amplified via PCR for 13 cycles, barcoded, and pooled into sets of 10-12 samples for sequencing on an Illumina HiSeq 2500. Illumina adapters and low-quality score (<20) bases were removed from the raw reads using TrimGalore! V.0.2.7 (67). Trimmed reads were mapped to the rhesus macaque genome (*MacaM* v7) using the STAR 2-pass method (68). Following quality control, we retained RNA-seq for 125 samples: 42 NC samples, 40 LPS samples, and 43 Gard samples.

Gene-level counts were obtained using STAR (69). Prior to RNA-seq data analysis, we first filtered out genes that were very lowly or not detectably expressed in our samples. Specifically, we removed genes that exhibited low median RPKM (≤ 2) in all three conditions, which resulted in a final set of read counts for 9,088 genes. We then normalized gene expression levels across samples using the TMM algorithm (weighted trimmed mean of M-values), implemented in the R package *edgeR* (70). Finally, we log-transformed the data using the *voom* function in R package *limma* (71, 72).

Gene expression levels can differ across social groups because of unknown environmental differences between groups. Additionally, samples from females living in the same social group were almost always collected at the same time (i.e., blood samples from all five members of the a group were drawn, cultured, shipped, and extracted in parallel). Hence, controlling for social group effects simultaneously controls for unmeasured variation between groups and most technical batch effects related to sample collection and processing. We also controlled for potential flow cell effects to take into account batch effects introduced at the sequencing stage.

To correct for batch effects, we implemented a mixed model to estimate the effects of social group (i.e., collection batch) and flow cell on gene expression, while taking into account dominance rank and other biological covariates of interest. To model past rank, we used Elo score evaluated a mean of 3.64 ± 1.06 s.d. months after females were introduced into their Phase I groups. Within each condition, we also quantile normalized past Elo scores to match the distribution of Elo scores for current rank, standardized the two Elo score variables and the two tissue composition variables to mean=0 and standard deviation=1, and mean-centered age. For each gene and each condition (NC, LPS, Gard) separately, we fit the following model:

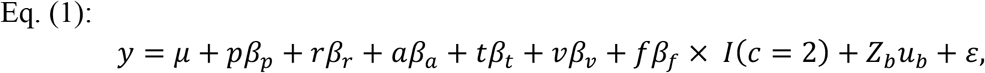

where the normalized expression level, *y*, is modeled as a function of an intercept,*μ*, and the fixed effects of past rank, *p* (*β_p_* is its coefficient); rank at the time of sampling, *r* (*β_r_* is its coefficient); age at the time of sampling, *a* (*β_a_* is its coefficient); the top two principal components of the normalized cell type proportions data, *t* and *v* (with coefficients *β_t_* and *β*_ν_, respectively); and flow cell, *f* (*β_f_* is its coefficient). For the flow cell effect, we only modeled an effect for the Gard condition (condition *c*=2) because Gard samples were sequenced on two flow cells; for LPS and NC conditions, all samples were sequenced on the same flow cell. Finally, we modeled a random effect, *u_b_*, of social group membership, which captures most sources of batch effects. *Z_b_* is an incidence matrix that assign samples to social groups. *ε* denotes model error.

We fit each model using the *lme* function in the R package *lme4* (73). To obtain batch-corrected values, we then subtracted the sample-specific estimate of the random effects of social group and flow cell from the original gene expression values. We note that this procedure only corrects for batch effects within, but not across conditions. The major source of batch effects across conditions arises from differences in the timing of library prep and sequencing for the Gard samples, relative to the NC and LPS samples (which were prepped and sequenced together). Because condition is completely collinear with this source of batch effects, we cannot correct for it in our data analysis. Importantly, this issue does not impact our estimation of rank effects *within* NC, LPS, or Gard data sets, which are the effects of primary interest for this study. It could, however, impact the identity and magnitude of our estimates for genes that are differentially expressed in response to Gard stimulation (i.e., when we compare NC to Gard data collected from the same individuals). Reassuringly, gene set enrichment analysis shows that genes we identified as responsive to Gard are strongly enriched for pathways involved in antiviral responses (e.g., response to cytokine: FDR-corrected p=2.4×10^−6^; type I interferon signaling pathway: FDR-corrected p=1.2×10^−5^; cellular response to interferon gamma: FDR-corrected p=3.1×10^−5^; *SI Appendix*, Dataset S2). This observation suggests that differences in expression between Gard and NC samples reflect a true biological response to the viral ligand and are not a technical artifact of our sequencing design.

### Statistical modeling of rank effects

To investigate rank effects on gene expression levels upon bacterial versus viral stimulation, and to quantify social history effects on these responses, we modeled batch-corrected gene expression levels using the following nested mixed model, with predictor variables quantile normalized (rank data), transformed to a standard normal (rank and tissue composition data), or mean-centered (age data) as described above:

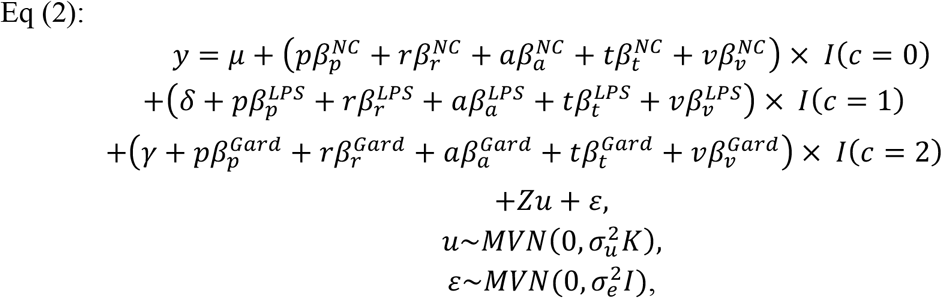

where *y* represents the normalized gene expression levels corrected for social group and flow cell effects, and data from all three conditions are modeled jointly. Here, gene expression levels are modeled as a function of an intercept, *μ;* the effects of LPS stimulation (*δ*) and Gard stimulation (*γ*); and the fixed effects of past rank, *p*; rank at the time of sampling (“current rank”), *r*; age, *a*; and the top two principal components of the normalized cell type proportions data, *t* and *v*. Fixed effects are nested within condition (control = 0; LPS = 1; Gard = 2), and *I* is an indicator variable for evaluating whether each sample was collected in the given condition. Coefficients are consistent with Eq. (1), but with an added superscript to denote estimates specific to a given treatment condition. We also model an *m* by 1 vector *u* as a random effects term to control for kinship and other sources of genetic structure. Here, *m* is the number of unique females in the analysis (*m*=45) and the *m* by *m* matrix *K* contains estimates of pairwise relatedness derived from a 45 × 54,165 genotype matrix, as described in (18). *Z* is an incidence matrix of 1’s and 0’s that maps samples to individuals in the random effects term. Residual errors are represented by *ε*, 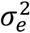 represents the environmental variance component (unstructured by genetic relatedness), *I* is the identity matrix, and MVN denotes the multivariate normal distribution.

To specifically estimate the effects of dominance rank on the response to LPS and Gard, we reformulated Eq. (2) to extract estimates for the interaction between rank and treatment condition, separately for past rank and current rank.

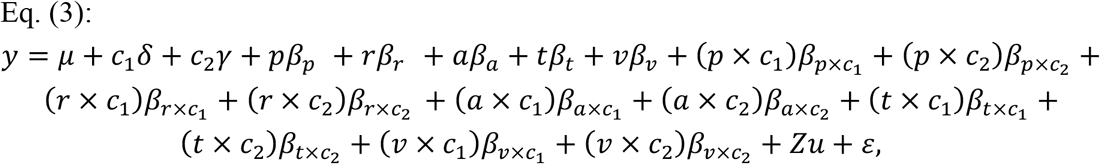

where the notation is consistent with Eq. (2), but we reparametrize the model to obtain directly each covariate’s interaction with treatment condition (i.e., the effects of each covariate on treatment response). These interactions can be interpreted as the effect of past rank, current rank, age, and tissue composition on the gene expression response to LPS or Gard challenge.

Finally, to explicitly model interaction effects between past rank and current rank, we introduced an additional term for each condition into Eq. (2):

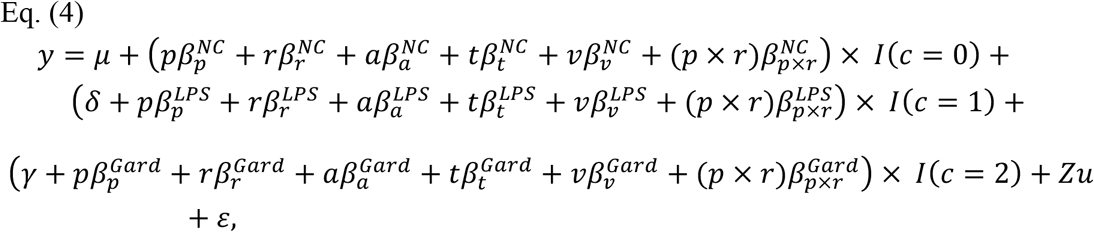

where notations are consistent with Eq. (2). Additional terms 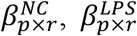, and 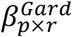 to represent condition-specific interactions between past and current rank.

We fit models 2 – 4 using the R package EMMREML (74). To assess significance after correcting for multiple hypothesis testing, we compared the distribution of p-values for each variable of interest to the distribution of p-values from permuted data. To generate permutations that preserved the structure of our data set (especially the strong correlation between Phase II social group membership and past rank in Phase I, which is a product of our experimental design: see Fig. 1A), we followed a two-step permutation procedure.

First, we blocked all explanatory variables (past rank, current rank, animal age, tissue composition variables) by social group in Phase II. We then permuted these blocks across social group labels (n=9 blocks and labels: one set for each social group), while keeping the gene expression data associated with the original social group label. Second, we shuffled the individual sets of explanatory variables randomly within each (now randomly re-assigned) social group. This two-step procedure maintains the correlation between Phase II social group and past rank observed in the original data because all co-housed females remain assigned to a shared social group label following permutation. However, it also randomizes the relationship between gene expression and the explanatory variables (see *SI Appendix*, Fig. S7 for a graphical depiction of this two-step strategy).

We performed permutations independently for each gene, conducted our complete set of statistical analyses (i.e., batch effect removal followed by differential expression analysis) on the permuted data, and treated the resulting p-value distributions as empirical null distributions to calculate false discovery rates, following a generalized version of the method described by Storey and Tibshirani (75).

To identify genes that significantly responded to Gard and LPS stimulation, which should not be affected by correlations between past rank and Phase II social group, we used the standard method for multiple testing correction implemented in the R package qvalue (76).

### Quantification of gene-specific plasticity

To quantify the plasticity of rank effects for each gene in each condition, we first estimated the total variance in gene expression levels explained by both past rank and current rank, 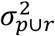. We then calculated the proportion of the total variance that could be accounted for by the effects of current rank in Phase II alone, 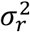. We define gene-specific plasticity, Θ, as the ratio of square roots of these variances (i.e., a ratio of standard deviations):

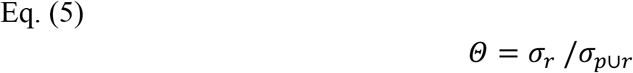

Since current rank and past rank are distributed following a standard normal (mean=0, s.d.=1) and almost perfectly uncorrelated (r=0.06, p=0.68), Θ can be approximated by the following expression:

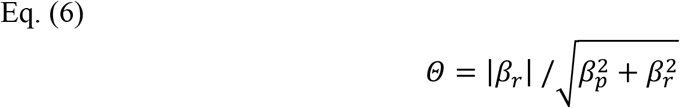

*Θ* therefore has an elementary geometric interpretation, which can be interpreted as the absolute value of the sine of the angle, *α*, created by the vector (*x, y*) = (*β_p_, β_r_*)and the x-axis (see *SI Appendix*, Fig. S7 for a schematic representation).

To make the distributions of *Θ* directly comparable across conditions, we focused our analyses on the top 1000 genes with the largest values of 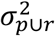 within each conditions (i.e., the genes most strongly affected by dominance rank, whether past, current, or both). We excluded the small set of genes for which past and current ranks were significant but directionally opposed (FDR=10%; <7.5% of all rank-associated genes in each condition). Importantly, the overall pattern--that social history effects are less apparent in immune-challenged conditions than in control conditions—is robust across a wide range of top rank-associated genes (*SI Appendix*, Fig. S6).

### Gene Ontology enrichment analysis

For all GO term enrichment analyses, we used the Cytoscape module ClueGO (77) (*SI Appendix*, Dataset S3). In all cases, we performed one-tailed Fisher’s Exact Tests for enrichment, and corrected for multiple tests using the Benjamini-Hochberg (B-H) method (78). To reduce the multiple testing space and account for the nested nature of GO terms, we analyzed only terms that fell between levels 3 and 8 of the GO tree for Biological Processes, included at least 5 genes in our data set, and for which at least 5% of the total number of genes belonging to the GO term were present in the test gene set. We report significant gene set enrichments for those GO categories that passed a 5% FDR for the response to LPS/Gard challenge and for current rank effects, and a 10% FDR for past rank effects.

### Transcription factor binding site enrichment analysis

To investigate whether binding sites for specific TFs were enriched in the promoter regions of rank-associated genes, we drew on chromatin accessibility data generated via ATAC-seq and reported previously in (18). In brief, these data were generated from 50,000 PBMCs obtained from three mid-ranking study subjects. The resulting libraries were sequenced on an Illumina NextSeq 500 and the reads mapped to the macaque genome and used to identify open chromatin peaks following the pipeline described in (18).

To identify likely TFBSs that overlapped with accessible chromatin upstream of rank-responsive genes (within 5 kb of the transcription start site), we scanned the macaque genome for matches to 1900 TRANSFAC and JASPAR-derived position weight matrices (PWMs: threshold log_2_ likelihood ratio(TFBS/background)=13) (79, 80). To reduce redundancy, we used bedtools (81) to perform hierarchical clustering of the PWMs based on the pair-wise Jaccard distances between their locations in the macaque genome. We then defined independent TF clusters at a dissimilarity threshold of 0.2. After filtering for clusters that rarely occurred in open chromatin regions upstream of genes, we obtained a final set of 460 motif clusters that we tested for enrichment near rank-associated genes.

## AUTHOR CONTRIBUTIONS

JS, MEW, JT, and LBB designed the study. NSM, NDS, TV, JK, and VM collected the data. JS and PLM analyzed the data. JT, LBB, and JS wrote the paper, with contributions and edits from all authors. JT, LBB, VM, and MEW provided funding support.

## ACKNOWLEDGMENTS

We thank J. Whitley, A. Tripp, N. Brutto, and J. Johnson for maintaining the study subjects and collecting behavioral data; A. Dumaine, V. Yotova and J. Brinkworth for experimental support; A. Bailey for access to wet lab facilities at YNPRC; M. Gutierrez for help with figures; and members of the Barreiro and Tung labs for helpful comments and discussion. This work was supported by NIH Grants R01-GM102562, R01-AG057235, P51-OD011132, K99/R00-AG051764, and T32-AG000139; NSF Grant SMA-1306134; the Canada Research Chairs Program 950-228993; NSERC Grant RGPIN/435917-2013; and North Carolina Biotechnology Center Grant 2016-IDC-1013. J.S. was supported by the Fonds de recherche du Québec–Nature et technologies, the Fonds de recherche du Québec–Santé, and the Canadian Institutes of Health Research Banting fellowship.

