## Supplemental Figures 1 to 7 for "Social history and exposure to pathogen signals modulate social status effects on gene regulation in rhesus macaques"

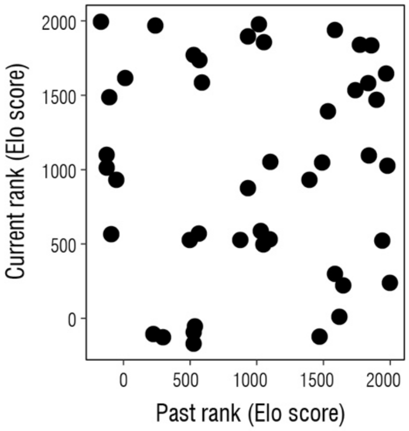


**Figure S1. No within-individual correlation between dominance ranks in Phase I and Phase II.** Our experimental paradigm produced completely uncorrelated Elo score estimates of current rank (at time of sampling) and past rank in Phase I (Pearson's r=0.06, p=0.68). This study design allowed us to separate the effects of current rank from rank social history. Each dot represents a single study subject (n = 45).

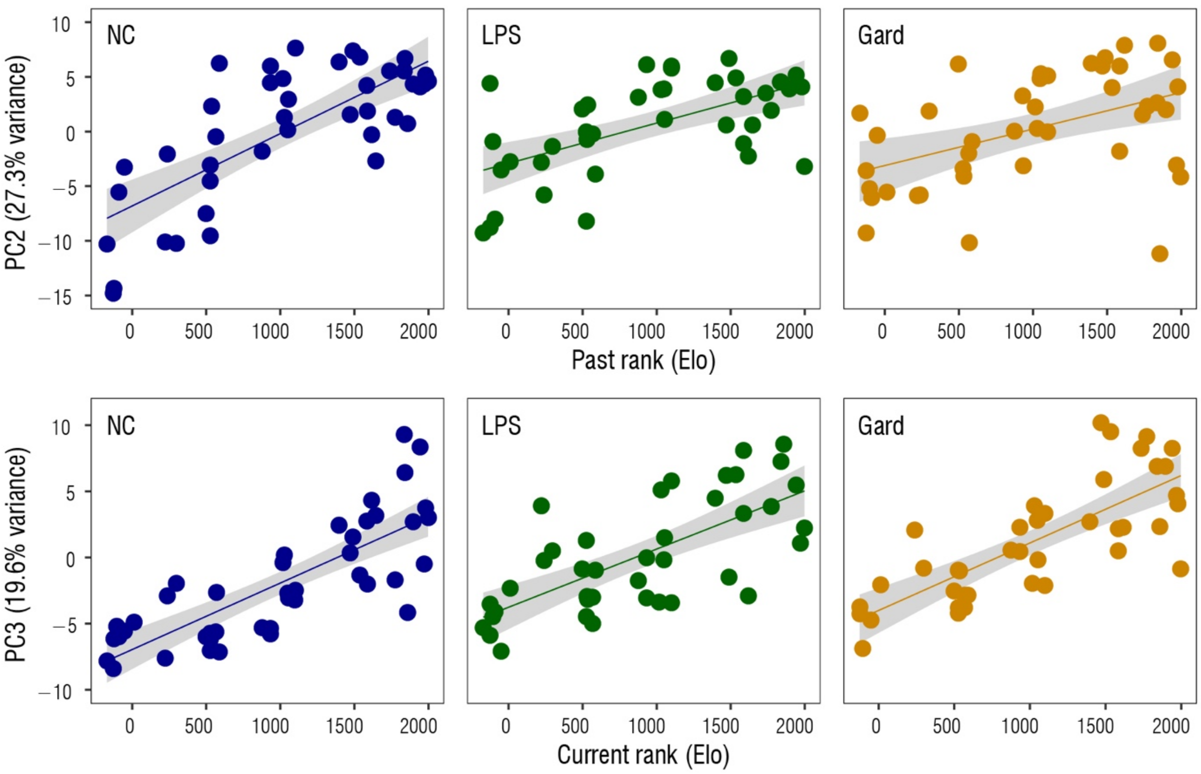


**Figure S2. Correlation between dominance rank (Elo score) and principal components from the PCA of data from all three conditions.** *(A)* Correlation between loading on PC3 (y-axis) and current rank (x-axis), in control (NC), LPS, and Gard conditions (Pearson's r = 0.75 (control); r = 0.75 (LPS), r = 0.69 (Gard), p<6.0 x $10^{-7}$). *(B)* Correlation between loading on PC2 (y-axis) and past rank in Phase I (x-axis), in control (NC), LPS, and Gard conditions (Pearson's r = -0.76, p=2.4 x $10^{-9}$; LPS: r = -0.58, p=8.0 x $10^{-5}$; Gard: r = -0.44, p=3.8 x $10^{-3}$).

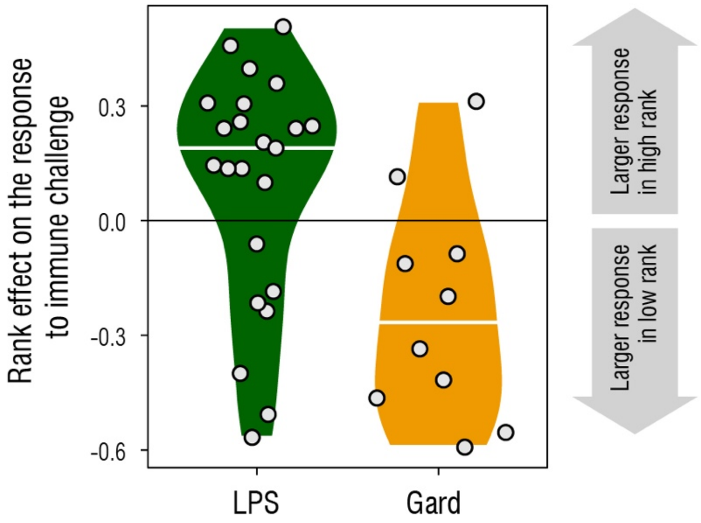


**Figure S3. Contrasting effects of dominance rank on the gene expression response of type I IFN-associated genes to LPS versus Gard.** Polarization of rank effects for genes in the Gene Ontology category "response to type I interferon" that are upregulated by LPS/Gard challenge. Positive values on the y-axis correspond to genes for which the gene expression response to LPS (left: green violin plot) or Gard (right: orange violin plot) is larger in high-ranking females; negative values correspond to genes for which the gene expression response to LPS or Gard is larger in low-ranking females. These results highlight the *change* in gene expression from unstimulated control conditions to LPS/Gard-stimulated conditions (in contrast to main text Figure 2C, which focuses on gene expression within LPS or Gard conditions). The distribution of rank effects for the response to LPS and Gard differ (Wilcoxon test: p=0.006); only genes that exhibited a relaxed evidence of association between dominance rank and the response to LPS/Gard (FDR<20%) are shown.

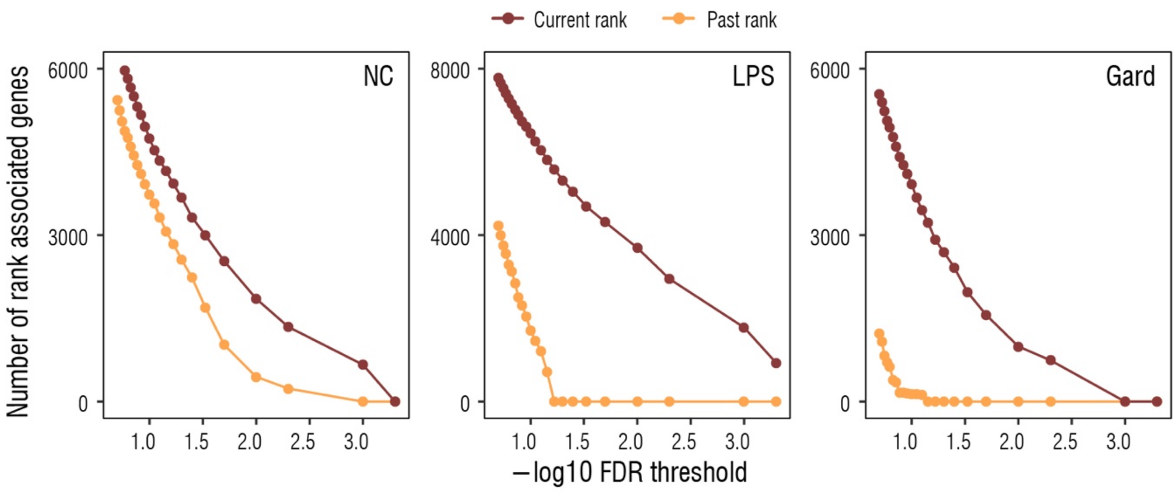


**Figure S4. Current rank effects and past rank effects across conditions and FDR thresholds.** Number of genes for which gene expression is explained by current rank in Phase II (brown lines) and past rank in Phase I (orange lines) at different FDR cutoffs (x-axis). Current rank effects are more frequent than past rank effects in all conditions, but past rank effects are more common in the unstimulated control samples than LPS- or Gard-stimulated samples regardless of FDR cutoff.

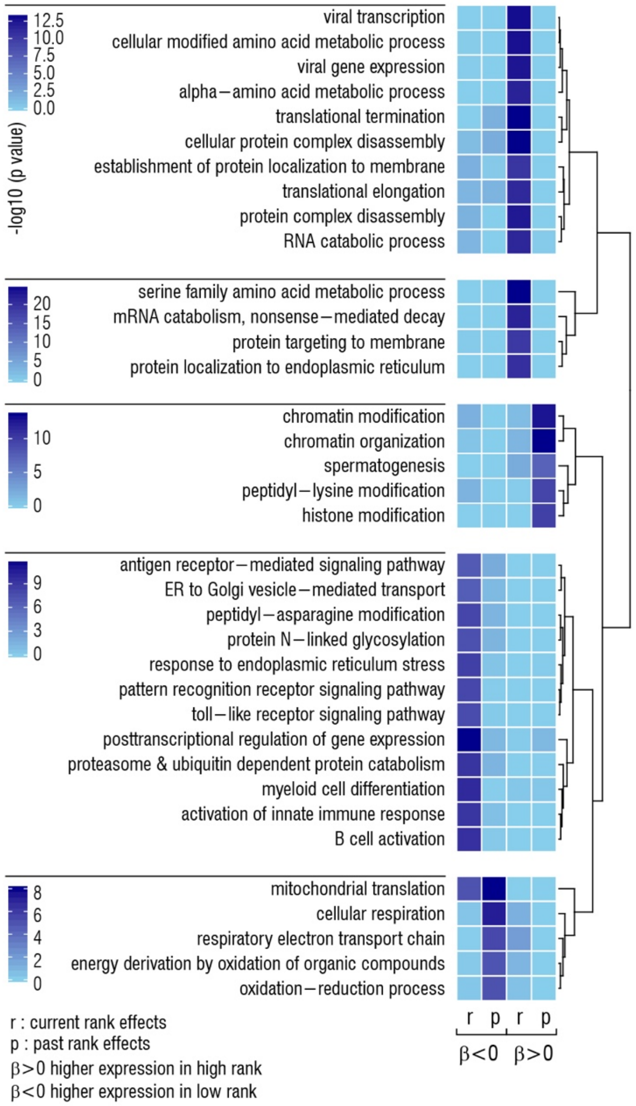


**Figure S5. GO terms enriched among current rank-responsive and past rank-responsive genes at baseline.** The heatmap shows a subset of significantly enriched Gene Ontology biological process terms, grouped by sets of related terms. Darker blue squares correspond to stronger statistical support for enrichment, and are ordered (left to right) for genes that are more highly expressed with (i) low current rank; (ii) low past rank; (iii) high current rank; and (iv) high past rank. Because we are less well-powered to detect past rank effects, rank-associated genes in this figure are defined based on an FDR of 5% for current rank and a relaxed cutoff of 10% for past rank effects. For a full list of GO terms enriched in each case,  see *SI Appendix*,  Dataset S2).

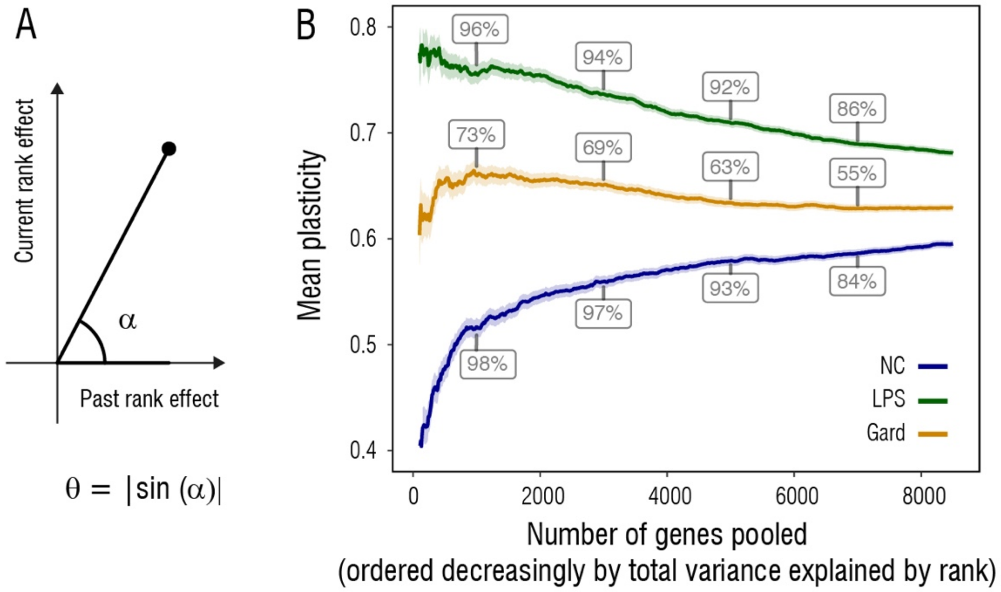


**Figure S6. Plasticity scores are lower in the unstimulated control samples than in the LPS or Gard conditions.** *(A)* The plasticity score Θ can be interpreted as the absolute value of the sine of the angle, $\alpha$, created by the vector x,y=(p,r)and the x-axis. Thus, Θ values range from 0 to 1, where values close to 1 imply a high degree of plasticity and little evidence of memory (i.e., strong effects of current rank and no effect of past rank), and values close to 0 imply a high degree of memory and little plasticity in response to changes in social status (i.e., strong effects of past rank and no effect of current rank). *(B)* Mean plasticity scores (y-axis) for each condition, calculated for the genes most affected by dominance rank. We ranked all genes in decreasing order by total variance in gene expression explained by the joint contributions of past and current rank, and calculated mean values of Θ for the top *x* genes, where *x* is shown on the x-axis. Boxed numbers show the percentage of genes included, at the corresponding x-axis threshold, that are significantly associated with current rank, past rank, or both (at 10% FDR). In the main text, we show distributions and summary statistics for the top 1000 genes; however, regardless of the number of genes included in these analysis, plasticity scores are systematically lower in baseline than in the LPS or Gard conditions.

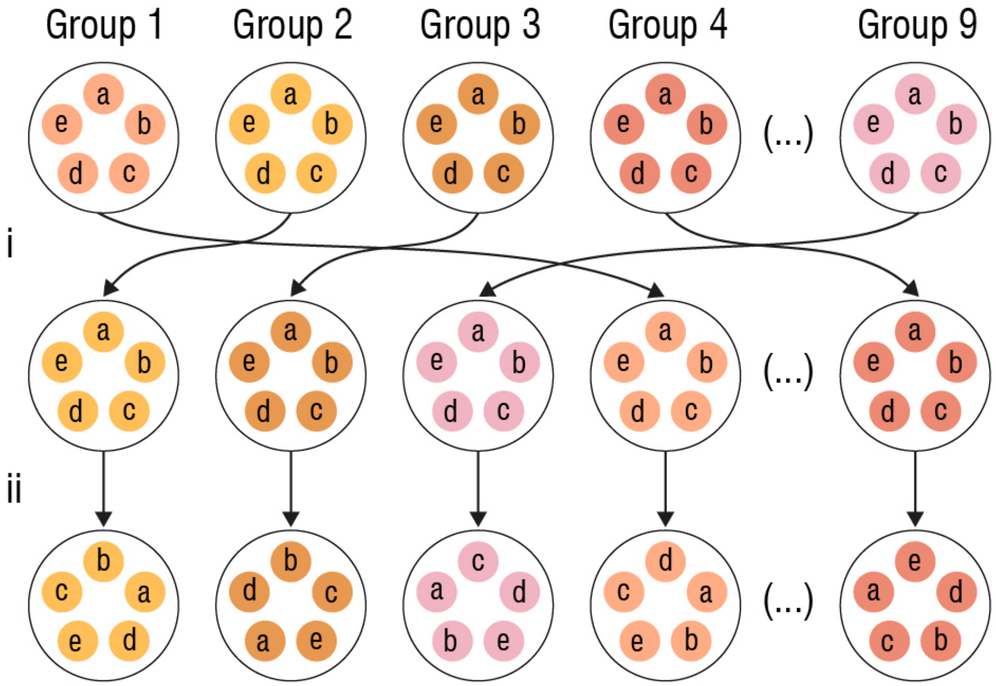


**Figure S7. Schematic representation of the permutation strategy for defining the empirical null p-value distribution for past rank effects.** To generate permutations that preserved the structure of our data set we followed a two-step permutation procedure. First (step *i*), we blocked all explanatory variables (past rank, current rank, animal age, tissue composition variables) by social group in Phase II. We then permuted these blocks across social group labels (n=9 blocks and labels: one set for each social group), while keeping the gene expression data associated with the original social group label. Second (step *ii*), we shuffled the individual sets of explanatory variables randomly within each (now randomly re-assigned) social group. This two-step procedure maintains the correlation between Phase II social group and past rank observed in the original data because all co-housed females remain assigned to a shared social group label following permutation. However, it also randomizes the relationship between gene expression and the explanatory variables.
